# Transient light-activated gene expression in Chinese hamster ovary cells

**DOI:** 10.1101/2020.10.22.351445

**Authors:** Shiaki A. Minami, Priya S. Shah

## Abstract

**Background:** Chinese hamster ovary (CHO) cells are widely used for industrial production of biopharmaceuticals. Many genetic, chemical, and environmental approaches have been developed to modulate cellular pathways to improve titers. However, these methods are often irreversible or have off-target effects. Development of techniques which are precise, tunable, and reversible will facilitate temporal regulation of target pathways to maximize titers. In this study, we investigate the use of optogenetics in CHO cells. The light-activated CRISPR-dCas9 effector (LACE) system was first transiently transfected to express eGFP in a light-inducible manner. Then, a stable system was tested using lentiviral transduction.

**Results:** Transient transfections resulted in increasing eGFP expression as a function of LED intensity, and activation for 48 hours yielded up to 4-fold increased eGFP expression compared to cells kept in the dark. Fluorescence decreased once the LACE system was deactivated, and a half-life of 14.9 hours was calculated, which is in agreement with values reported in the literature. In cells stably expressing the LACE system, eGFP expression was confirmed, but there was no significant increase in expression following light activation.

**Conclusions:** Taken together, these results suggest that optogenetics can regulate CHO cell cultures, but development of stable cell lines requires optimized expression levels of the LACE components to maintain high dynamic range.

## Background

Chinese hamster ovary (CHO) cells are a major industrial workhorse for mammalian glycoprotein production. Improving protein yields through regulation of gene expression is a major focus of CHO cell engineering [1]. Reversible and scalable control of gene expression would represent a major step forward in CHO cell engineering, yet mature methods typically rely on constitutive regulation or on chemical induction that cannot be easily reversed unless the media is replaced [2]. For instance, sodium butyrate is a highly utilized chemical in industrial CHO cell cultures to increase specific productivity [3]. However, the increase in specific productivity is often accompanied by a decrease in cell viability [4]. Removing sodium butyrate towards the end of the culture duration may improve viability while retaining the benefits of increased specific productivity [5], but media replacement is economically disadvantageous in large bioreactors. A system in which stimuli are easily removed would allow greater control over the entire culture period, which is crucial given the dynamic nature of production cultures.

Recently developed systems, such as the light-activated CRISPR-dCas9 effector (LACE), enable regulation of gene expression with a tunable and easily reversed mechanism [6]. Briefly, the LACE system relies on light-activated dimerization of CRY2 and CIBN. CIBN is fused to a CRISPR/dCas9 system, where the guide RNA (gRNA) localizes the CIBN-dCas9-CIBN complex to a minimal CMV promoter. CRY2 undergoes a conformational change and dimerizes with CIBN upon excitation with 450 nm light. CRY2 fusion to the viral transactivation domain VP64 enables transcriptional activation upon light-activated dimerization. Removal of the light reverses the dimerization to turn off transcription. This is a modular system in which light can be used to regulate the expression of any transgene of interest (**Fig 1**).

**Figure 1.**
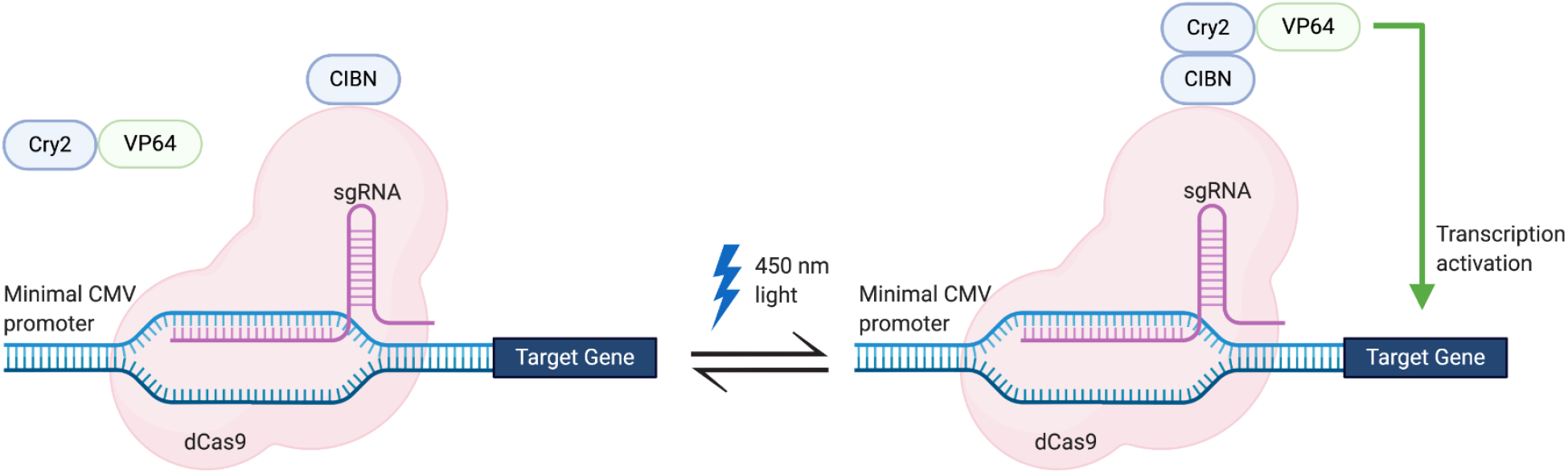
Schematic of LACE system. The LACE system relies on light-activated dimerization of CRY2 and CIBN (blue). CIBN is fused to dCas9 and targeted to the minimal CMV promoter using a synthetic gRNA sequence. CRY2 is fused to VP64 so that it activates transcription following light activation and dimerization with CIBN.

Here, we use this previously described LACE system to control gene expression in CHO cells. We show that transient light-activated expression of enhanced green fluorescent protein (eGFP) is possible through transient transfection of a four-plasmid system, and this system is tunable. However, only a small percentage of cells received adequate amounts of all four plasmids, and the transient nature of the system limits its utility for CHO cell engineering applications. We therefore tested stable expression relying on lentiviral transduction. This system did not result in light-activated expression of eGFP, suggesting additional modifications are required for stable light-activated gene expression.

## Results

### LACE activity in CHO cells using transient transfection

We first tested the LACE system in CHO-DG44 (DG44) cells using transient transfection and eGFP as a reporter for expression. Since the minimal CMV promoter can result in leaky transcription, we determined the background level of eGFP expression by transfecting only the eGFP plasmid (minCMV-eGFP). Background MFI of eGFP signal was slightly lower than that of the full four-plasmid LACE system without light activation. Following light activation for 24 hours, eGFP expression was measured by flow cytometry. We observed an approximately 4-fold increase in mean fluorescence intensity (MFI) for the population of cells that were gated positive for eGFP expression (**Fig 2A**).

**Figure 2.**
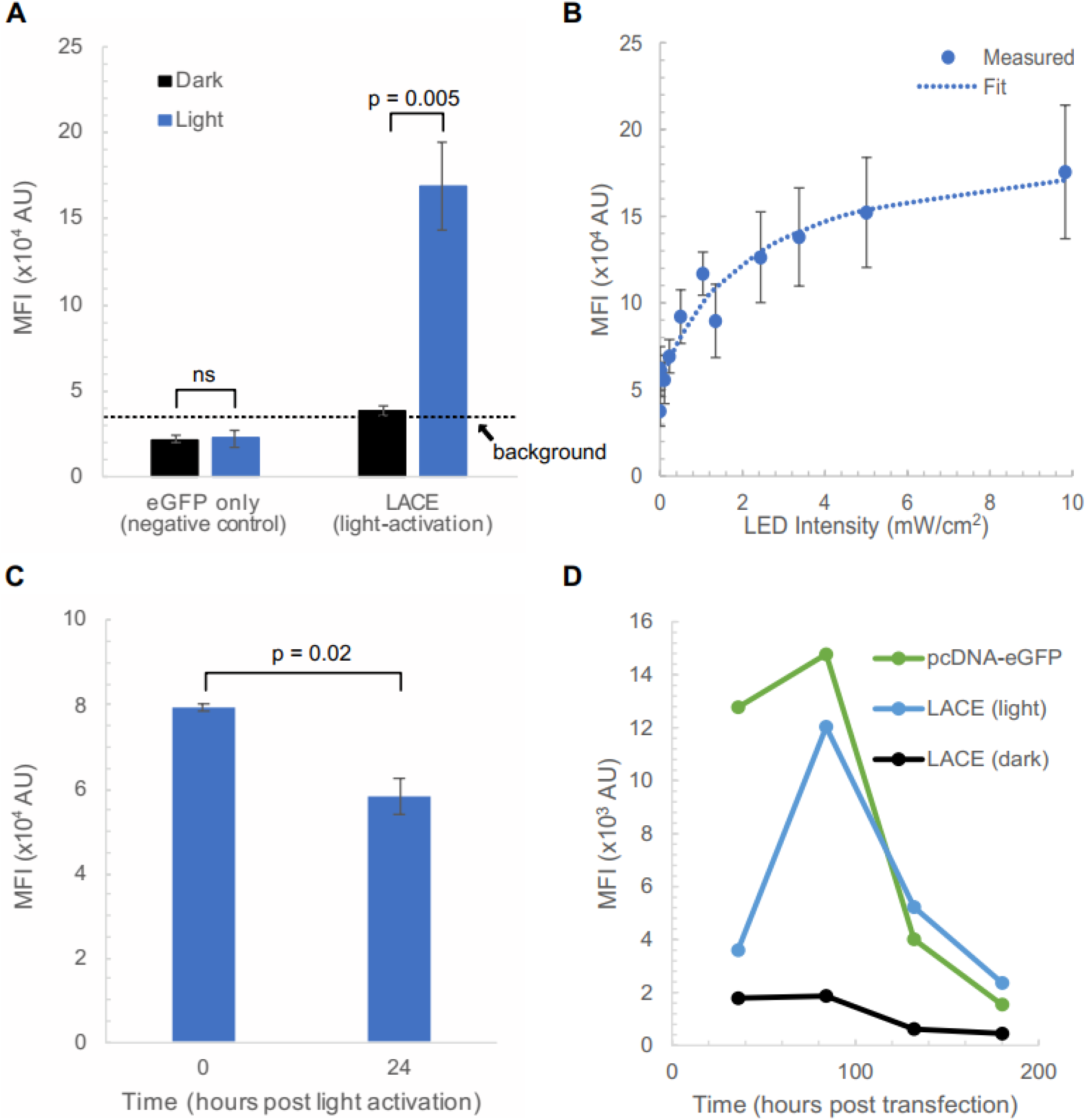
Transient light-active gene expression in CHO cells. The LACE system was used to express eGFP in CHO cells in a light-activated manner. **(A)** DG44 cells were transfected with LACE plasmids. Mean fluorescence intensity (MFI) of eGFP signal was measured by flow cytometry with (blue) and without (black) 24 hours of light activation with 450 nm light using 1 s pulses at 0.067 Hz pulse frequency and 9.8 mW/cm^2^ intensity. minCMV-eGFP (negative control) represents the background fluorescence of cells (dashed line). **(B)** Tunability of eGFP expression was measured by altering LED intensity. MFI of eGFP signal from DG44 cells was measured by flow cytometry following 24 hours of light activation at the indicated intensity. A fit of the data was performed assuming Michaelis-Menten kinetics. **(C)** Reversibility of eGFP expression was measured by monitoring eGFP signal from DG44 cells after turning off light activation. MFI was measured by flow cytometry 0 and 24 hours after light activation was terminated. **(D)** CHO-K1 cells were transfected and with the indicated plasmids. Cells transfected with LACE plasmids with either exposed to light or kept in the dark. MFI of eGFP signal was measured at the indicated time. Data from **(A)** and **(B)** represent the mean values of five and three independent experiments, respectively. Data from (**C**) represent the mean values of three technical replicates in an experiment. Error bars represent standard error of the mean. Background fluorescence is indicated with a dashed line and the upper limit of the 95% confidence interval for negative control samples. P values were calculated with a paired two-tailed T-test.

Once we confirmed that the LACE system functions in CHO cells, we sought to test tunability, reversibility, and duration of light-activation. We tested whether eGFP expression could be tuned in CHO cells using light intensity. We varied light intensity while keeping pulse width and frequency constant. We observed saturating behavior in which more than 80% of maximum MFI of eGFP signal was achieved with only 50% of the maximum light intensity possible (**Fig 2B**). We also showed that the system is reversible in CHO cells. MFI of eGFP fluorescence decreases when light activation is terminated (**Fig 2C**). Using an exponential decay fit, the half-life of eGFP intensity was calculated to be ~14.9 hours. This is within the range of commonly reported values in the literature [7, 8]. Together, our results show that the LACE system is tunable and reversible in CHO cells. We next tested how long light activation could be maintained in CHO cells using transient transfection. CHO-K1 cells were transfected with the LACE plasmids and subjected to light activation for the indicated times post-transfection or kept in the dark. MFI of eGFP signal was measured by flow cytometry. The decay in MFI of eGFP signal for the LACE system following light activation (light) mirrored the decay by constitutively active eGFP (pCDNA-eGFP). We observed a peak MFI of eGFP signal at 84 hours post-transfection, which was a 6-fold increase compared to control cells kept in the dark. While the signal was still moderately high and well above control cells kept in the dark at 132 hours post-transfection, at 180 hours post-transfection the signal was indistinguishable from control cells kept in the dark (**Fig 2D**). Together, our results show that the LACE system is tunable and reversible in CHO cells, with light-activated expression viable for up to 5 days post-transfection.

### LACE activity in CHO cells using stable lentiviral transduction

Many applications in CHO cells require the generation of stable cell lines and cultures that last more than one week [9, 10]. Transient systems are becoming more popular, but given the decay of the LACE signal following transfection (**Fig 2D**), LACE activity would only be accessible in the first five days post-transfection. Moreover, transient transfection of the LACE plasmids would result in only a small percentage of the cells receiving all four plasmids required for LACE activity. For example, even with a transfection efficiency as high as 90%, only 66% (0.9^4^) of cells would receive four plasmids of an equimolar mixture. In these cases, stable expression of LACE components would be desirable. We therefore cloned the LACE components into a lentiviral expression system using Gibson Assembly. Each component was introduced to DG44 by serial transduction. We then split cells into two samples with one sample cultured in the dark and the other sample activated with light for 24 hours. While we did not observe significant light-activation, a small population of cells (~0.1%) had increased fluorescence following light activation (**Fig 3A**). To test whether this result was cell-line dependent, we tested transient transfection and serial transfection in CHO-K1 cells. Very similar results were obtained for CHO-K1, with robust transient light-activated expression of eGFP (data not shown), but low signal-to-noise for stably integrated cells (**Fig 3B**). We hypothesized that the population of light-responsive cells was very small because only a small percentage of them received and appropriately expressed all four plasmids using serial transduction. We therefore sorted single CHO-K1 cells in the top 0.1% of fluorescence intensity following light activation to further characterize their response to light. Clones were allowed to recover and expand for more than two weeks before analysis. Surprisingly, light activation of the sorted clones resulted in the same MFI of eGFP signal as the clones without light activation. This almost perfect 1:1 relationship between dark and light for the stably-transduced clones stands in contrast to the approximately 4-fold change in MFI for the transient system and indicates that the sorted clones are not light-responsive (**Fig 3C**).

**Figure 3.**
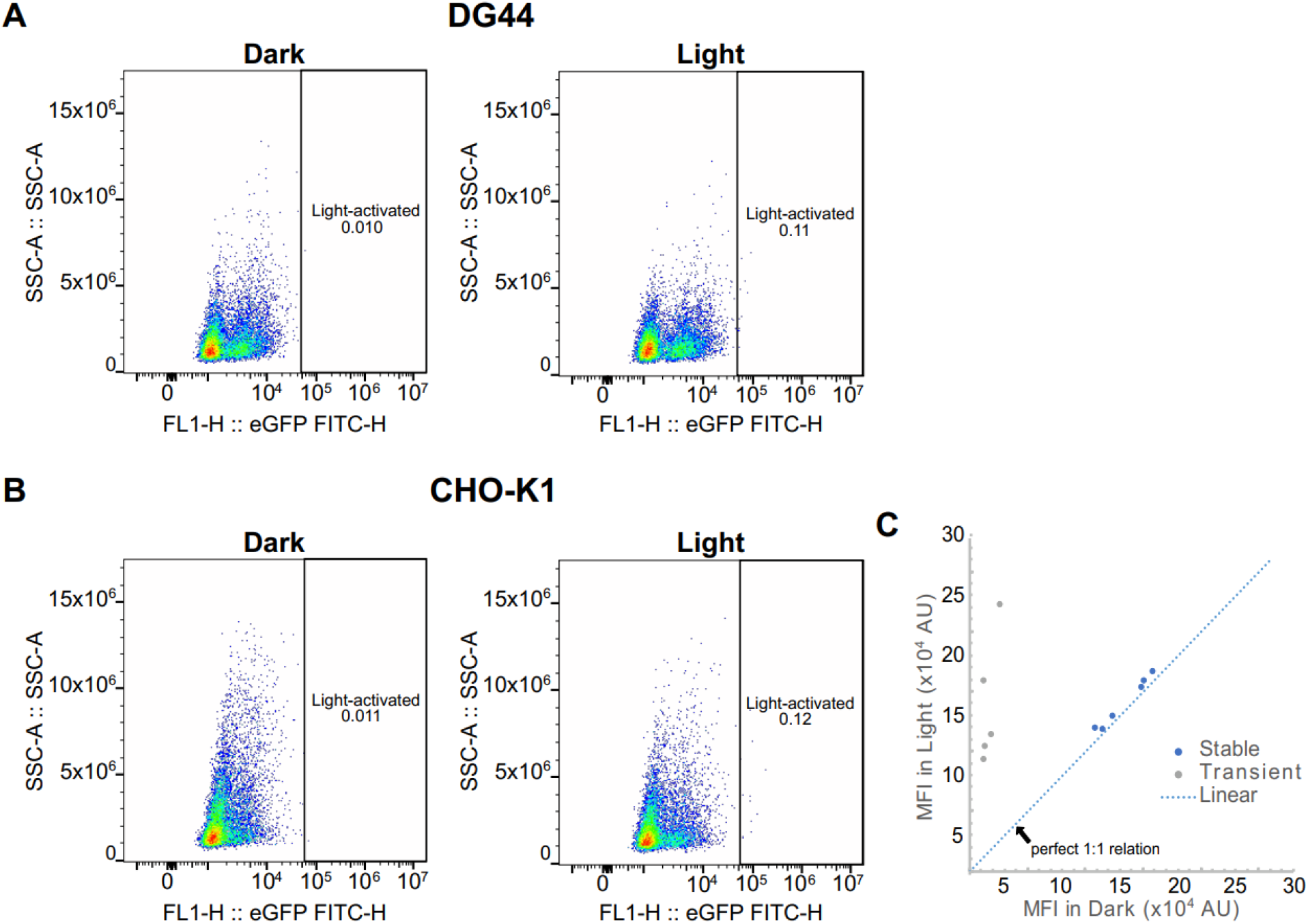
Lentiviral transduction does not result in stable light-activated gene expression. **(A)** DG44 and **(B)** CHO-K1 cells serially transduced with lentiviral vectors encoding the LACE system were incubated in the dark or with light activation for 24 hours. Fluorescence intensity of cells was measured by flow cytometry. **(C)** Cells in the top 0.1% of fluorescence from **(B)** were single-cell sorted and expanded. The surviving clonal populations were incubated in the dark or with light activation for 24 hours. Fluorescence intensity of cells was measured by flow cytometry. Transient transfection data from **Fig 2A** is included as a reference for robust light activation.

Given the absence of light-activation in the lentiviral system, we sought to determine if each of the lentiviral constructs retained activity. We therefore tested the lentiviral plasmids in a transient transfection setting for responsiveness to light activation. Each of the lentiviral constructs except for the eGFP-encoding construct was tested in combination with the transient transfection-based LACE plasmids. When only one lentiviral construct was included, we observed between 2- and 6-fold increases in MFI of eGFP signal following light activation. The lentiviral construct encoding the gRNA resulted in a light-activated MFI similar to that of the signal of cells left in the dark for other combinations, suggesting that this construct may not be functional. However, in experiments in which two lentiviral constructs were used in tandem with two transient transfection-based LACE plasmids, we still observed robust 4-fold light activation. Interestingly, MFI of eGFP signal for cells kept in the dark increased as we added more lentiviral constructs to the transient transfection. This increase in background fluorescence was not always accompanied with proportional increases in signal following light activation, suggesting that diminishing signal-to-noise ratio for the lentiviral system limits its utility (**Fig 4A**). We were not able to test the lentiviral construct encoding eGFP in the transient transfection system because of its intrinsic fluorescence observed during transient transfection scenarios such as lentiviral packaging (**Fig 4B**).

**Figure 4.**
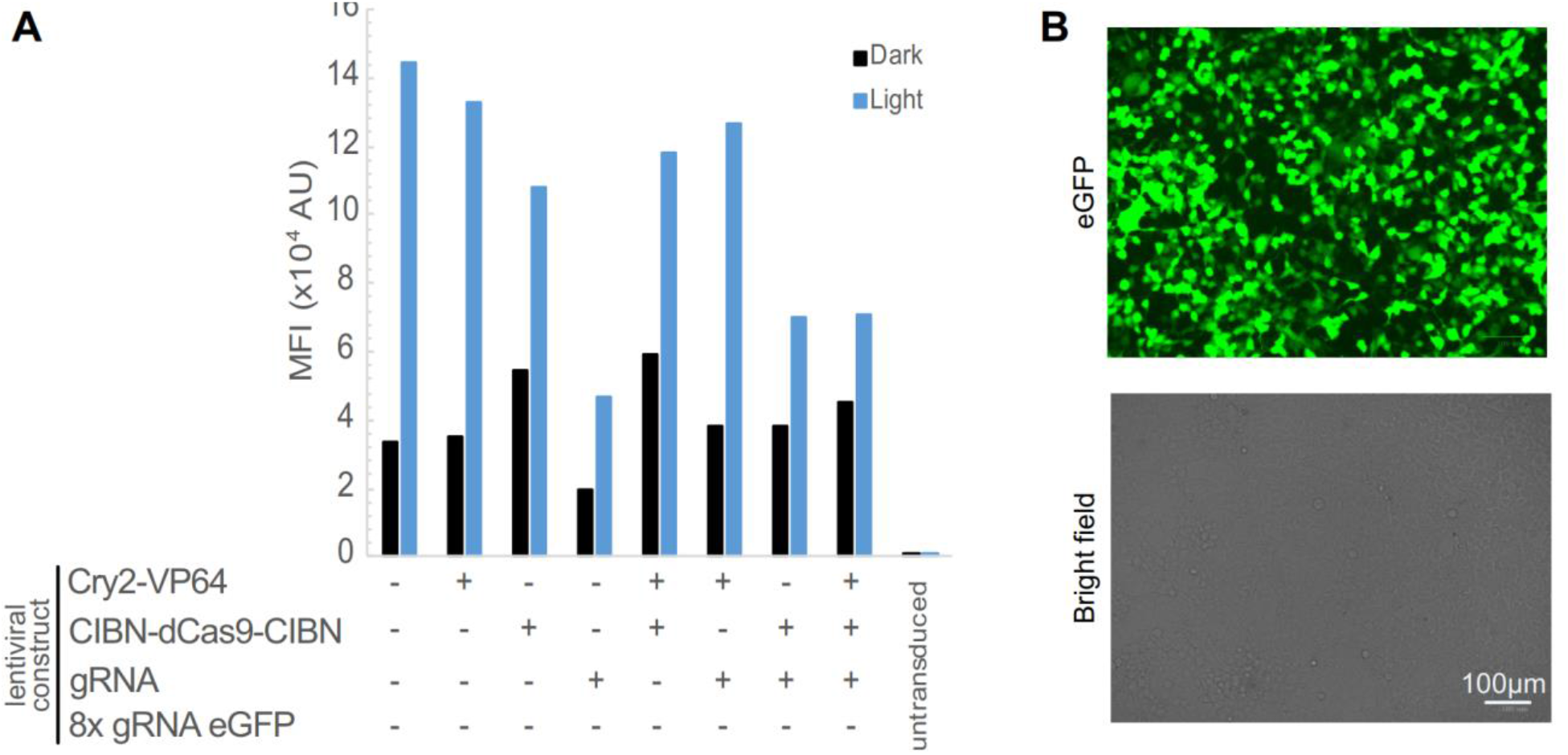
Lentiviral constructs function in a transient setting. **(A)** The indicated combinations of transient and lentiviral LACE plasmids were transiently transfected into CHO cells and incubated in the dark or with light activation. MFI of eGFP signal was measured by flow cytometry after 24 hours of light activation. The 8xgRNA eGFP lentiviral construct was not tested because of its intrinsic eGFP expression in a transient setting (see **(B)**). **(B)** eGFP expression of the 8x gRNA eGFP lentiviral construct. HEK 293T cells were transfected with the 8x gRNA eGFP lentiviral construct along with lentiviral packaging plasmids. eGFP and bright field images were taken 24 hours post-transfection.

## Discussion

Here we test the LACE system for light-activated gene expression in CHO cells. We find that transient transfection of LACE plasmids results in tunable and reversible light-activated gene expression that can last approximately five days (**Fig 2**). The deactivation of the LACE system is dependent on the half-life of the protein being expressed because the time scale for LACE component interaction is much smaller than the time scale for protein degradation. Thus, if desired, the rate of reversion of the system into the dark state can be improved by using destabilized proteins. High transfection efficiencies are necessary (> 95%) because this system relies on co-transfection of four separate plasmids. Moreover, this transient system would not allow for applications involving stable cell lines, or experiments spanning more than one week. We therefore tested the LACE system in lentiviral constructs. We found that the lentiviral LACE system had low signal-to-noise that limited its application (**Figs 3 and 4**). We allowed clones to recover for at least two weeks to allow them to relax back into the dark state, but it is still possible that the previous light activation required to sort individual clones created an epigenetic memory that increases the background fluorescence of these clones, even in a prolonged dark state [11]. Identifying cells that include all four components of the LACE system using antibiotic selection markers and fewer constructs could be used to circumvent this potential technical barrier.

It is also possible that the lentiviral LACE system exhibited low signal-to-noise because of increased intrinsic background eGFP expression from the minimal CMV promoter in the lentiviral setting. Our observations of high eGFP expression from the 8x gRNA eGFP lentiviral construct in a transient transfection setting (**Fig 4B**) support this hypothesis. In this case, a lentiviral system may not be a viable option as a stable LACE system. While less efficient than lentiviral transduction, transient transfection and stable cell line generation via antibiotic selection could circumvent the need for lentiviral transduction. Alternatively, incorporating mammalian origins of replication into all LACE plasmids could allow for episomal plasmid maintenance following transient transfection [12].

## Conclusions

In summary, transient light-activated gene expression in CHO cells is tunable and reversible using the LACE system. While stable expression of LACE components would expand their utility, lentiviral transduction of LACE components resulted in a low signal-to-noise ratio. Other approaches to stable expression of LACE components, such as antibiotic selection of transfected LACE plasmids, could overcome this technical challenge.

## Methods

### Plasmids

pcDNA3.1-CRY2FL-VP64, pcDNA3.1-CibN-dCas9-CibN, pGL3-Basic-8x-gRNA-eGFP, and gRNA-eGFP-Reporter were gifted from Charles Gersbach (Addgene plasmid # 60554, 60553, 60718, and 60719, respectively). These plasmids were used directly for transient transfection. Lentiviral vectors were constructed from PCR amplification of the LACE components. CRY2FL-VP64 was amplified with primers 1 and 2, CibN-dCas9-CibN with primers 3 and 4, and gRNA-eGFP-Reporter with primers 5 and 6. A synthesized eGFP-IRES-mTagBFP2 fragment was PCR-amplified with primers 7 and 8. mTagBFP2 was introduced so that LACE system activation could be monitored once eGFP was replaced with another gene of interest. PCR-amplified fragments were cloned using Gibson Assembly into pFUGW vectors, gifted from Isei Tanida [13] (Addgene plasmid # 61460), which were digested with PacI (New England Biolabs) and EcoRI-HF (New England Biolabs).

**Table 1.**
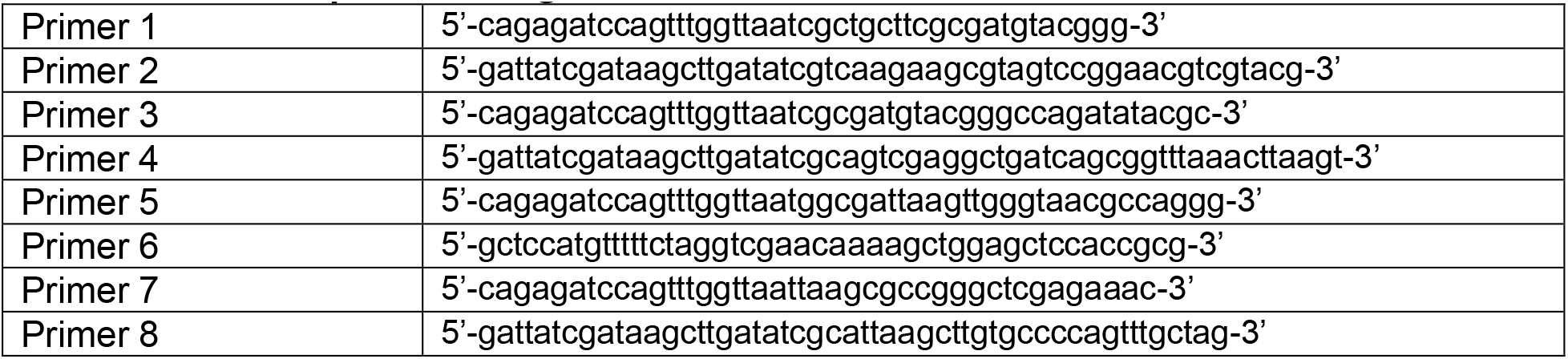
Primer sequences for generation of lentiviral vectors

### Cells

CHO-K1 cells (ATCC) were maintained in IMDM (Gibco) supplemented with 10% FBS (Gibco) and 1% penicillin/streptomycin (Lonza). CHO-DG44 cells [14] were maintained in IMDM supplemented with 10% FBS, 1% penicillin/streptomycin, 13.6 mg/L hypoxanthine (Alfa Aesar), and 3.9 mg/L thymidine (Sigma). HEK293T cells (ATCC) were maintained in DMEM (Gibco) supplemented with 10% FBS and 1% penicillin/streptomycin. Cells were cultured in a humidified incubator at 37 °C and 5% CO_2_. Cell density and viability were measured using trypan blue and an automated cell counter (TC20, Bio-Rad).

### Transient transfection

At 24 hours before transfection, cells were seeded in 12-well plates at a density 0.12 x 10^6^ cells per well. To introduce the LACE system, cells were transfected with 206 ng pcDNA3.1-CRY2FL-VP64, 294 ng pcDNA3.1-CibN-dCas9-CibN, 100 ng pGL3-Basic-8x-gRNA-eGFP, and 400 ng gRNA-eGFP-Reporter. Transfections were performed using PolyJet DNA *In Vitro* Transfection Reagent (SignaGen Laboratories) with a PolyJet/DNA ratio of 2.

### Lentiviral packaging and transduction

Lentivirus was packaged in HEK293T cells using sodium phosphate co-transfection of 3.5 ug transgene, 1.8 μg gag/pol-, 1.25 μg vsv-g-, and 0.5 μg rsv-rev-expressing plasmids [15]. Medium exchange was performed 6-12 post-transfection. Medium containing lentivirus was harvested 48-60 hours post-transfection via centrifugation at 1000 rpm for 10 minutes and filtration through 0.45 μm syringe filters. Packaged lentivirus was stored at −80 °C until further use.

For transduction, cells were plated on 12-well plates at a density 0.12 x 10^6^ cells per well and transduced serially with the LACE system components. Cell culture medium was replaced by 250 μL medium containing CRY2FL-VP64 lentivirus and 250 μL fresh medium. After 1 hour, 500 μL fresh medium was added. The following day, cells were passaged back to a density of 0.12 x 10^6^ cells per well to prepare for the next transduction. CIBN-dCas9-CIBN, gRNA-eGFP-Reporter, and 8x-gRNA-eGFP-IRES-BFP were then individually transduced in this manner.

### Light activation

24 hours post-transfection, cells were activated via 8mm LEDs (465 nm, L.C. LED). LEDs were mounted on a breadboard, and intensities and pulse frequencies were regulated using an Arduino Uno microcontroller. For all experiments, cells were illuminated with 1-second pulses of light, with a pulse frequency of 0.067 Hz. Unless otherwise specified, LED intensities were 9.8 mW/cm^2^. LED intensities were measured using an optical power meter (1931-C, Newport). Samples in the dark were wrapped in aluminum foil to ensure isolation from light.

### Flow cytometry

Cells were trypsinized (Gibco) and resuspended in culture medium. Following centrifugation at 300 rcf for 5 minutes at 4 °C, supernatants were discarded, and cell pellets were resuspended in cold DPBS containing 1% FBS. A 488 nm laser was used for all flow cytometry analysis.

BD FACSCanto II was used to analyze LACE system activity over 180 hours. CytoFLEX S Flow Cytometer was used for all other analytical cytometry experiments. Gating for eGFP-positive cells were performed such that >99.9% of untransfected cells were excluded.

Astrios MoFlo EQ was used to sort single cells. Following the activation of serially transduced cells, cells were activated with 465 nm light for 24 hours. Cells with eGFP fluorescence intensities in the top 0.1% of the population were sorted into 96-well plates containing cell culture medium. Surviving colonies were scaled up to 12-well cultures before light-activation and flow cytometry analysis.

### Fitting eGFP MFI vs LED intensity and determining the half-life of eGFP

Production of eGFP was modeled assuming a saturation production rate model with first-order degradation and a constant term for low expression that is observed in the dark:

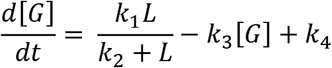

[*G*] is the mean fluorescence intensity of eGFP, *L* is the intensity of LED light, *k*_1_ and *k*_2_ are constants describing the saturation, *k*_3_ is the degradation rate constant, and *k*_4_ is the constant for constitutive expression. To fit the parameter scan for eGFP MFI vs. LED intensity, the steady-state solution was used:

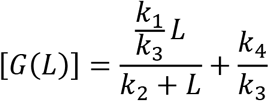

The equation was fit using fitnlm in MATLAB, yielding 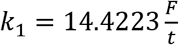, 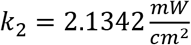, *k*_3_ = 1.0073 *t*^−1^, and 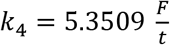. F is fluorescence intensity of eGFP in arbitrary units.

To calculate protein half-life, the time-dependent equation was solved after setting *L =* 0 to apply the effect of turning the LEDs off:

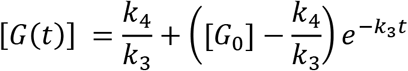

[*G*_0_] is the fluorescence intensity of eGFP at the time the LEDs are turned off. Fitting this expression, it was found that 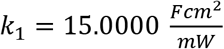, *k*_2_ = 1.0000 *F*, *k*_3_ = 1.2581 *t*^−1^, and 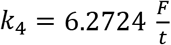. The degradation rate constant, *k*_3_, is similar between both fits performed. The half-life was then calculated with first order degradation kinetics:

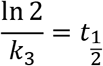

Using *k_3_* values from the LED intensity and reversibility experiments, half-life of eGFP was determined to be an average of 14.9 hours.

## List of Abbreviations

(CHO): Chinese hamster ovary
(DG44): CHO-DG44
(LACE): Light activated CRISPR-dCas9 effector
(CRISPR): Clustered Regularly Interspaced Short Palindromic Repeats
(gRNA): guide RNA
(eGFP): enhanced green fluorescent protein
(mTagBFP2): monomeric Tag Blue Fluorescent Protein
(MFI): mean fluorescence intensity

## Declarations

### Ethics approval and consent to participate

No ethics approval or consent was required for this research.

### Consent for publication

No consent for publication was required for this research.

### Availability of data and materials

The processed datasets supporting the conclusions in this article are included within the article. All raw data used to generate plots and materials will be made available upon request.

### Authors’ contributions

SAM and PSS conceived of the project, designed experiments and wrote the manuscript. SAM performed all experiments. Both authors read and approved the final manuscript.

## Acknowledgements

We thank Kush B. Patel for assistance with cloning and Nitin S. Beesabathuni for feedback on project development and the manuscript. We also thank Dr. Florian Krammer for providing CHO-DG44 cells.

